# Importance of vitamin D in critically ill children with subgroup analyses of sepsis and respiratory tract infections: a systematic review and meta-analysis

**DOI:** 10.1101/390476

**Authors:** Margarita Cariolou, Meghan A. Cupp, Evangelos Evangelou, Ioanna Tzoulaki, Antonio J. Berlanga-Taylor

## Abstract

**Background:** Critical care and sepsis remain high priority concerns in children. Observational studies report high prevalence of vitamin D deficiency and present mixed results regarding the correlation between vitamin D status and adverse outcomes. Associations between deficiency and mortality, particularly in children with sepsis, remain unclear. We performed a systematic review and meta-analysis to address this uncertainty.

**Methods:** PubMed, OVID and Google Scholar were searched for observational studies in critically ill children. We obtained pooled prevalence estimates for vitamin D deficiency and odds ratios for the association of mortality in critically ill children treated in intensive care units, with subgroup analysis for those with sepsis and those with respiratory tract infections. Meta-regression and sensitivity analyses were used to investigate heterogeneity.

**Findings:** Forty-eight studies were included. Total sample size was 7,199, with 1,679 (23%) children acting as controls in case-control studies. Of 5,520 critically ill children, 2,664 (48%) were vitamin D deficient (< 50 nmol/L). Results of the random effects model demonstrated a pooled prevalence of vitamin D deficiency of 54·9% (95% CI 48·0-61·6, I^2^=95·0%, 95% CI 94·0-95·8, p < 0·0001). In subgroup analysis of children with sepsis (16 studies, 788 total individuals) we observed higher prevalence of deficiency (63·8%, 95% CI 49·9-75·7, I^2^=90·5%, 95% CI 86·2-93·5%, p < 0·0001). In patients admitted for respiratory tract infections (24 studies, 1,683 total individuals), prevalence was 49·9% (95% CI 37·6-62·2; I^2^= 93·9%, 95% CI 92·1-95·3, p < 0·0001). Only one identified study assessed vitamin D levels in sepsis and mortality. A meta-regression model with all available variables (year of publication, total study sample size, quality score, study design, country group and clinical setting) explained 37·52% of I^2^(F = 5·1119, p = 0·0005) with clinical setting and country groups being significant predictors for prevalence.

Meta-analysis of mortality (18 studies, 2,463 total individuals) showed an increased risk of death in vitamin D deficient critically ill children both with random (OR 1·81, 95% CI 1·24-2·64, p-value = 0·002) and fixed effects (OR 1·72, 95% CI 1·27-2·33, p= 0·0005) models with low heterogeneity (I^2^= 25·7%, 95% CI 0·0-58·0, p = 0·153) and low evidence of publication bias (p = 0·084, Egger’s test). There were insufficient studies to perform meta-analyses for sepsis and respiratory tract infection related mortality.

**Interpretation:** Circulating vitamin D deficiency is common amongst critically ill children, particularly in those with sepsis. Our results suggest that vitamin D deficiency in critically ill children is associated with increased mortality. Clinical trials, studies with larger sample sizes and standardized approaches are needed to further assess associations between circulating levels of vitamin D and mortality and other outcomes in the paediatric population.

**Funding:** Medical Research Council UK

**Registration:** PROSPERO (CRD42016050638)

**Copyright:** Open access article under terms of CC BY

**Research in context:** *Evidence before this study:* Vitamin D deficiency is common worldwide and has been associated to numerous diseases in observational studies. The extent of deficiency and relevance to mortality in children receiving acute and intensive care is unclear and only recently has gained more attention. We searched PubMed, OVID, Google Scholar and the Cochrane Library from inception up until 5^th^November 2017 without language restrictions. Search terms used across these databases included: “critical care”, “acute care”, “vitamin D”, “pediatric”, “child”, “neonate”, “toddler”, “intensivecareunit”, “sepsis” and “septic shock” (full search terms are listed in the appendix). Most (81·3%) included studies were published between 2014 and 2017. We did not identify randomised clinical trials assessing the value of vitamin D supplementation in this population. We did not find sufficient studies to perform meta-analyses for mortality from sepsis or respiratory tract infection.

*Added value of this study:* Our systematic review and meta-analysis provides an in-depth assessment of the magnitude and relevance of vitamin D circulating levels in paediatric acute and critically ill patients with pre-specified sub-group analyses. We found that studies were highly heterogeneous across a number of important study variables including clinical setting, patient age groups, sample size, geographic location, case definitions, study quality, study design, biomarker thresholds and assay measurements. Pooled estimates of prevalence of vitamin D deficiency were overall extremely high, showing that around half of patients in general and acute paediatric care are deficient. Estimates were similar for respiratory tract infections but higher in sepsis, with overlapping confidence intervals across all outcomes. Individual study estimates were highly variable however. We analysed this heterogeneity using meta-regression and identified clinical setting and country of study as important contributors, likely indicating that patient age and environmental exposure to vitamin D, amongst other likely important factors, are key determinants and should be adequately assessed and reported. Pooled estimates for mortality outcomes showed a clear increased risk with lower vitamin D levels, despite the variation in study characteristics. We were unable to assess the importance of vitamin D levels in sepsis and respiratory tract infections due to the small number of studies investigating these outcomes.

*Implications of the available evidence:* Vitamin D deficiency in acute and critical care settings is common and associated with increased mortality in paediatric patients. Our review highlights the heterogeneous nature of the study population however and emphasizes the need for adequate power and control of confounding factors in future work. Few studies have investigated specific diseases such as sepsis and respiratory tract infections in relation to vitamin D despite their high prevalence, social and economic costs. Understanding the causal nature and therapeutic value of vitamin D in paediatric critical care remain key areas for investigation.

## Introduction

Vitamin D is an essential nutrient^1, 2^ representing a group of fat soluble secosteroids with key endocrine functions.^3^ It is synthesized in the skin upon sunlight exposure^4^ while dietary sources, such as oily fish, egg yolk, certain fungi and supplements, are usually secondary sources. Vitamin D is critical in bone metabolism^5^ and calcium homeostasis,^6^ as well as acting as an important regulator in extra-skeletal metabolic processes,^7^ cardiovascular and immune systems.^8^ Many observational and laboratory studies have observed the anti-inflammatory properties of vitamin D,^9^ including direct regulation of endogenous anti-microbial peptide production.^10^

It is therefore crucial for humans to have sufficient vitamin D levels to maintain bone health and possibly improve response to infection.^6, 11, 12^ Infants and children are especially dependent on vitamin D to achieve healthy bone development and growth.^13, 14^ Well-known functional outcomes of adequate vitamin D levels in children include rickets prevention, higher bone mineral content and reduced bone fracture rates.^5, 14^ In otherwise healthy children in the United States, the reported prevalence of vitamin D deficiency (25OHD levels of < 25 nmol/L) ranges from 9 to 18%.^15^ The Endocrine Society Clinical Practice Guidelines and the Institute of Medicine (IOM) suggest that vitamin D levels less than 50 nmol/L (20 ng/mL) reflect a deficient state.^4, 16^

Studies in adults reflect a high prevalence of vitamin D deficiency both in general intensive care unit (ICU) and sepsis patients and strongly suggest an association between low vitamin D and poor clinical outcomes, including increased mortality, particularly in those suffering from sepsis.^2, 17^ Recent clinical trials of vitamin D supplementation in adults appear promising in both general critical care^18, 19^ and sepsis.^20^

Sepsis remains a challenging clinical entity with high social and economic costs.^21^ Each year there are approximately 123,000 sepsis cases and around 37,000 deaths in England alone.^22^ Recent reports show an increased prevalence of paediatric sepsis,^23^ likely a reflection of an increased population with chronic comorbidities, higher rates of opportunistic infections and multidrug resistant organisms.^24^ Respiratory tract infections account for a large proportion of underlying diagnoses in acute and critical care conditions^24, 25^ but remain understudied.^26^

The magnitude and relevance of vitamin D deficiency in children receiving acute care is not clear. Several recent studies have addressed these questions with mixed results. We sought to summarise the evidence regarding the implications of vitamin D deficiency and its prevalence in general ICU, respiratory tract infection and sepsis patients in the paediatric population. We carried out a systematic review and meta-analysis of circulating vitamin D levels to assess the prevalence of vitamin D deficiency (≤ 50 nmol/L) and its association with mortality in these conditions.

## Methods

We planned and conducted our systematic review and meta-analysis according to the PRISMA guidelines^27^ and since we did not include randomized controlled trials we reported following the Meta-Analysis of Observational Studies in Epidemiology (MOOSE) guidelines.^28^

### Search strategy and selection criteria

Our population of interest consists of paediatric patients with acute conditions and/or treated in ICU or emergency units for acute conditions whose vitamin D status was assessed prior to or during admission. We included published cross sectional, case-control and cohort studies that measured circulating 25(OH) D levels and either reported prevalence, odds ratios (OR) or data to enable calculation. Studies were excluded if they were reviews, case reports, surveys, commentaries, replies, not original contributions, experimental *in vitro* or if they recruited patients who were not treated in emergency, neonatal intensive care units (NICUs), paediatric intensive care units (PICUs) or for acute conditions. Studies were also excluded if they only enrolled vitamin D deficient patients, investigated healthy populations only or did not measure circulating 25(OH) D levels as an indicator of vitamin D status. When we identified more than one publication utilising the same cohort, we included the publication which shared our review’s objective to investigate vitamin D levels and prevalence of deficiency.

For purposes of our review, we classified vitamin D deficiency as being less than 50 nmol/L (equivalent to 20 ng/mL) as suggested by the IOM.^16^ Different age categories were used to designate patients as “children” in the studies reviewed. We therefore included all “children” as defined by each treating facility and this included “neonates”, “infants”, “toddlers”, “children” and “adolescents”.

We searched PubMed, OVID, Google Scholar and the Cochrane Library from inception up until 5^th^ November 2017, with no language restrictions. Search terms used across these databases included: “critical care”, “vitamin D”, “pediatric”, “child”, “neonate”, “toddler”, “intensive care unit”, “sepsis” and “septic shock”. Search terms used in OVID and PubMed are listed in the *Additional Tables 1A and 1B*. Literature searches were performed by two investigators independently and included initial screening of titles and abstracts, followed by full text screening. Any disagreements for study eligibility were resolved by discussion between the authors. Reference lists of the selected papers, including reviews, were also checked for relevant titles. Abstracts of relevant titles were then assessed for eligibility. A data extraction form was designed a priori.

### Study quality assessment

The quality of each included study was assessed using the Newcastle-Ottawa Scale (NOS) for cohort, case-control and cross-sectional study designs.^29^ We classified studies as low (1-3), medium (4-6) or high quality (7-9) for purposes of sensitivity analysis (*Additional Table 2 A, B* and *C*).

### Prevalence and mortality outcomes

In the majority of studies (n = 36), prevalence of vitamin D deficiency was extracted as reported with a threshold of ≤ 50 nmol/L. If prevalence was not reported directly, it was calculated using data provided in each study (cases ≤ 50 nmol/L / total number of study participants, (*Additional Table 3 A* and *B*). Extracted or calculated prevalence values were then combined in a meta-analysis. For mortality, we calculated unadjusted odd ratios (OR) as:

OR = (vitamin D deficient patients who died * vitamin D non-deficient patients who did not die)/ (vitamin D deficient patients who did not die * vitamin D non-deficient patients who died)

We had sufficient information to calculate ORs < 50 nmol/L for 36 studies (75%). For the 12 studies with insufficient information, we used the lower cut-off values reported as a conservative approximation (*Additional Table 4*). We converted 25(OH) D values using: nmol/L = ng/mL * 2·496.

## Data analysis

We obtained proportions of vitamin D deficiency with 95% confidence intervals (CI) using the Clopper-Pearson method^30^ in R. We used a random effects model^31^ to account for the variation observed within and between studies due to the different ages and acute conditions in the populations considered. For mortality, we also obtained pooled proportions and pooled ORs with fixed effect model for sensitivity analysis and to avoid false conclusions that could result from small-study effects.^32^

We investigated possible sources of heterogeneity using sensitivity and subgroup analyses. The I^2^ statistic was used to estimate the percentage of total variation across studies which can be attributed to heterogeneity. A Q value of < 0·05 was considered significant and an I^2^ statistic greater or equal to 75% indicated a high level of variation due to heterogeneity.^33, 34^ We used Egger’s regression test to present results for publication bias and funnel plot asymmetry^35^ and generated funnel plots for visual assessment and screen for evidence of publication bias.

To further assess heterogeneity, we utilised meta-regression to identify predictor variables that could explain variation in study prevalence estimates. We used restricted maximum likelihood (REML) estimations in the model to account for residual heterogeneity^36^ and the Knapp-Hartung method to adjust confidence intervals and test statistics. This method estimates between study variance using a t-distribution, rather than a z-distribution, yielding a more conservative inference.^37^ We tested the following continuous predictors: year of study publication, total sample size and quality score. Categorical variables included study setting (PICU, NICU), study design (case-control, cross-sectional and cohort) and country group by geographic region and economic development (group 1, group 2, and group 3) and were dummy coded.

We used R version 3·5·0 and Microsoft Excel 2010 for analyses and data collection. The R packages “meta”^38^ and “metafor”^39^ were used for analyses. Only results of the random effects model are reported for prevalence due to the expected heterogeneity between populations being considered. Our protocol is registered in PROSPERO (CRD42016050638).

### Role of the funding source

The study received funding from the UK Medical Research Council. The funders had no role in data collection, analysis, interpretation or writing of the report. All authors had access to the data in the study.

## Results

### Screening and study characteristics

After title and abstract screening, we identified 2,890 potentially relevant studies (*Figure 1*) and eighty-five full text articles were assessed for eligibility. Rationale for study exclusion included: studies including adults, study populations other than critically ill children or with acute conditions, studies of circulating vitamin D levels and deficiency in healthy children or in children with chronic conditions. Four studies^40-43^ were excluded due to insufficient data reporting (*Additional Table 5*). We also excluded three studies^44-46^ that used the same cohort of children and included a single study to represent the cohort.^47^ Ultimately, 48 studies met criteria for inclusion (*Additional Table 6*).

**Figure 1.**
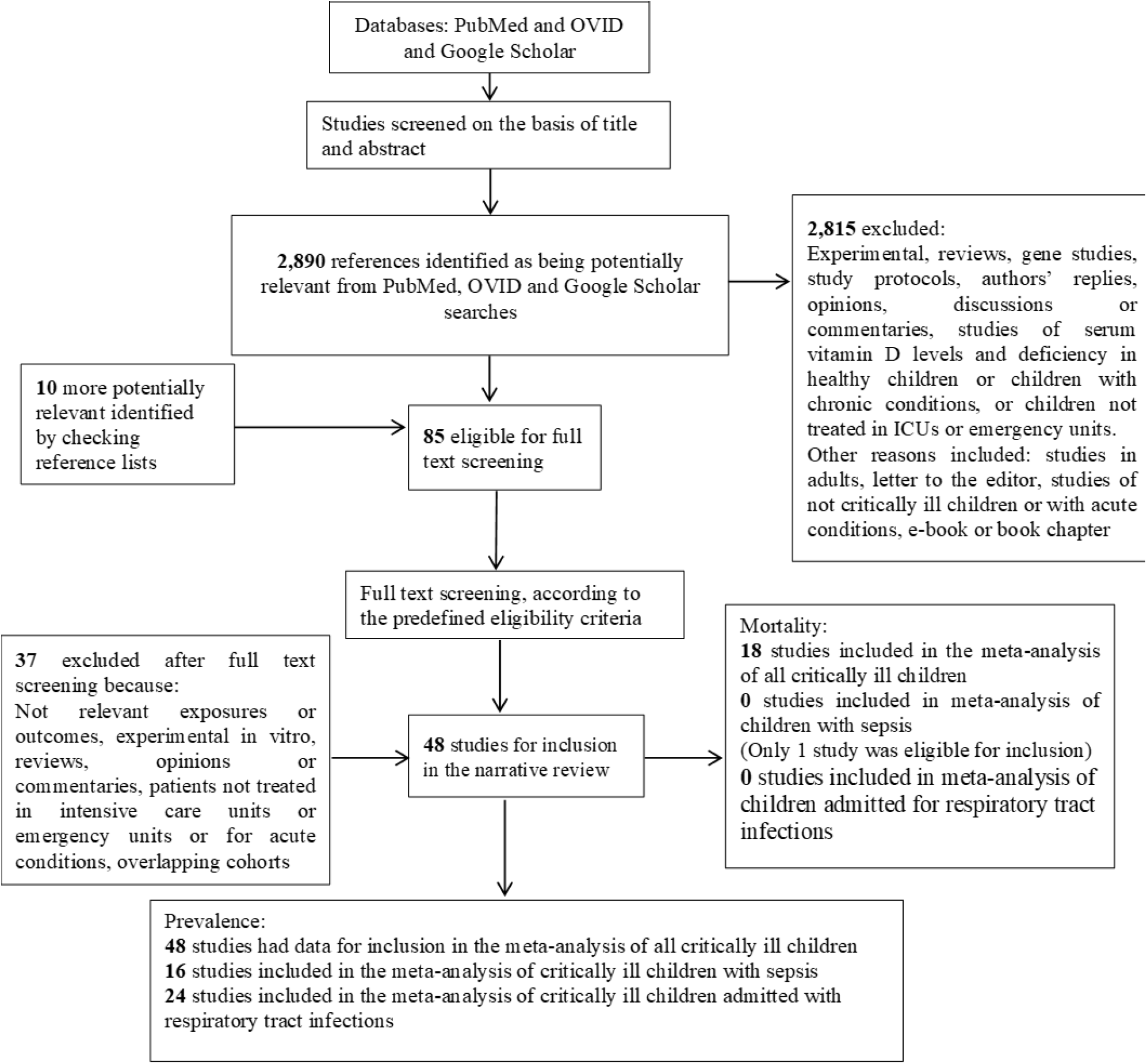
Flow chart of study selection process.

The primary objective of the majority of studies was to determine circulating vitamin D concentration (“status”) in children and/or prevalence of vitamin D deficiency. Secondary objectives included investigation of associations between deficiency of circulating vitamin D and various outcomes, such as hospital mortality length of stay, requirement of ventilation and/or illness severity (*Additional Table 7*).

All included studies reported vitamin D measurement assay methods used (*Additional Table 8*) and stated that samples were collected and analysed within the first 24 hours of hospital admission. Studies reported ethical approval and consent for participation from parents or guardians (*Additional Table 9*). Included studies were published between 2004 and 2017, with the majority (n = 39, 81·3%) published between 2014 and 2017 (*Additional Table 6*). In total, 5,520 children were hospitalized in paediatric or neonatal intensive care units or emergency units. Sample sizes of critically ill children ranged from 25^48^ to 511.^49^ In 16 studies the total number of cases was greater than 100.

Studies originated from 15 countries, with the majority from India^8, 50-58^ (n = 10) or Turkey^48, 59-64^ (n = 7) (*Additional Table 6*). All were of medium or high quality (NOS score median 6·5, range 4-8). The score range for cohort studies was 6 to 8 (n = 20), for case-control studies 5 to 8 (n = 24) and for cross sectional 4 to 6 (n = 4). Studies used a broad range of ages to classify patients as “children”. Six studies (12·5%)^48, 60, 62-65^ included only neonates. In two^60, 65^ of these six studies, neonates were preterm. The largest age range was seen in the study of Ayulo et al 2014, which included individuals between 1 and 21 years of age (*Additional Table 10*).

All studies included both female and male participants. For mortality, four of the 18 studies (22%) carried out multivariate regression analysis with adjustment for confounders. The remaining studies presented results using a variety of methods, including Spearman’s correlation analysis, chi-square or Fisher’s exact tests or descriptive statistics.

### Prevalence of vitamin D deficiency

We included 48 studies representing a total of 5,220 critically ill children. Of these, 2,664 (48%) were classified as vitamin D deficient (< 50 nmol/L). Prevalence of deficiency ranged from 5%^66^ to 95%^54^ (*Additional Table 11*). Sample sizes ranged from 25 to 511, with a median of 82 individuals (*Additional Table 12*). Using a random effects model, the pooled prevalence estimate of vitamin D deficiency was 54·9% (95% CI 48·0-61·6) with a high proportion of variation attributed to heterogeneity (I^2^ = 95·0%, 95% CI 94·0-95·8, p < 0·0001) (*Figure 2*) and evidence of funnel plot asymmetry (p = 0·015, Egger’s test) (*Table 1* and *Additional Figure 1*).

**Table 1.** Pooled estimates of vitamin D deficiency in critically ill children and critically ill children with sepsis and those with respiratory tract infections

| Patient category | Number of studies<br>(Total number of<br>individuals;<br>number of deficient<br>individuals) | Pooled proportion % (95% CI) | | Heterogeneity<br>($I^2$)<br>% (95% CI) | Q value,<br>d.f.<br>p-value | Eggers<br>p-value |
| --- | --- | --- | --- | --- | --- | --- |
|  |  | Random<br>effects | Fixed<br>effects |  |  |  |
| Critically ill children<br>(includes those with<br>sepsis) | 48 (5,520; 2,664) | 54.9<br>(48.0-61.6) | 46.8<br>(45.4-48.3) | 95.0<br>(94.0-95.8) | 931.46,<br>57,<br>< 0.0001 | 0.015 |
| Critically ill children<br>(only those with sepsis) | 16 (788; 499) | 63.8<br>(49.9-75.7) | 62.6<br>(58.6-66.5) | 90.5<br>(86.2-93.5) | 157.99,<br>15,<br>< 0.0001 | 0.828 |
| Only those admitted<br>with respiratory tract<br>infections | 24 (1,683; 778) | 49.9<br>(37.6-62.2) | 43.2<br>(40.4-46.1) | 93.9<br>(92.1-95.3) | 378.7,<br>23,<br>< 0.0001 | 0.217 |

CI = confidence intervals; $I^2$ = heterogeneity; df = degrees of freedom. Vitamin D deficiency defined in our study as < 50 nmol/L (20 ng/mL). $I^2$ statistic used to estimate heterogeneity between pooled studies: $I^2 \geq 75\%$ was considered high heterogeneity.

**Figure 2.**
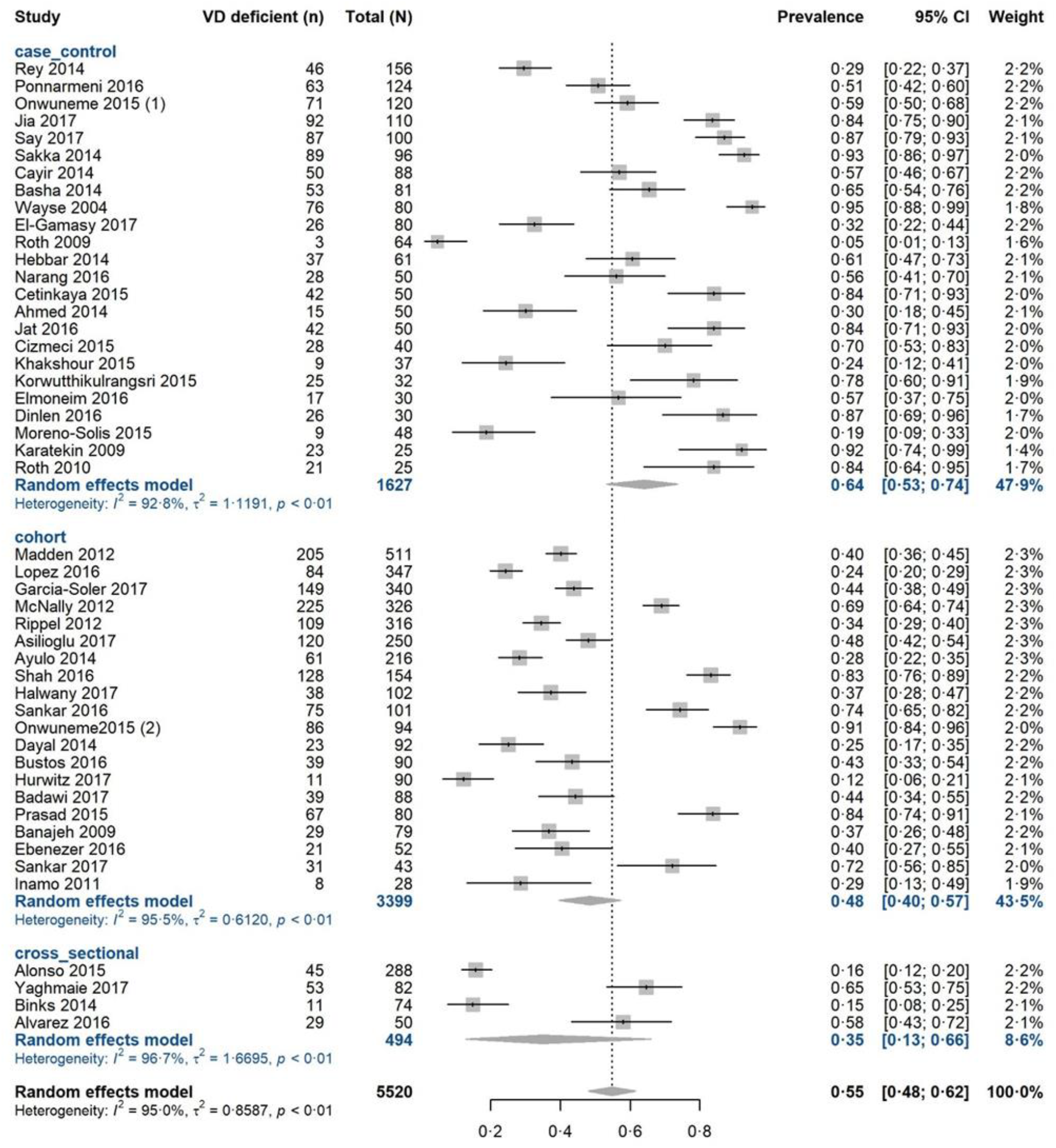
Pooled prevalence estimate for vitamin D deficiency in critically ill children by study design. Forest plot shows results from the random effects model. Each diamond represents the pooled proportion of vitamin D deficiency for each of the subgroups (case-control, cohort, cross-sectional study designs). The diamond at the bottom represents the overall pooled proportion of all the 48 studies together. Each square shows the prevalence estimate of each study and the horizontal line across each square represents the 95% confidence interval (CI) of the prevalence estimate.

### Sensitivity analysis for prevalence

We did not detect material differences in prevalence after exclusion of the 12 studies which did not directly report prevalence < 50 nmol/L (53·1%, 95% CI 45·6-60·4; I^2^ = 95·1%, 95% CI 93·9-96·0, p < 0·0001) (*Additional Table 13*).

When examining results by sample size (defining “large” as ≥ 100 and “small” as < 100), we found that the 16^8, 47, 49-51, 59, 60, 67-75^ studies with larger sample size included 3,561 total individuals and gave a prevalence estimate of 50·8% (95% CI 40·5-61·1; I^2^ = 96·9%, 95% CI 95·9-97·6, p < 0·0001). The remaining 32 studies with “smaller” sample sizes included 1,959 total children and estimated pooled prevalence as 57·2% (95% 47·3-66·7; I^2^ = 92·7, 95% CI 90·7-94·3, p < 0·0001) (*Additional Table 13*).

We also conducted analysis by study design. Cohort studies (n = 20) yielded a prevalence estimate of 48·4% (95% CI 39·7-57·3; I^2^= 95·5%, 95% CI 94·1-96·5, p < 0·0001). In case-control studies (n = 24) the estimate was 64·1% (95% CI 53·2-73·6; I^2^= 92·8%, 95% CI 90·5-94·6, p < 0·0001) and in cross-sectional (n = 4) 34·8% (95% CI 12·8-66·0; I^2^= 96·7%, 95% CI 94·0-98·2, p <0·0001) (*Additional Table 13*, *Figure 2*).

We assessed whether studies’ country of origin influenced results. Studies in India gave an estimate of 69·5% (95% CI 53·0-81·5; I^2^= 93·6%, 95% CI 90·2-95·8, p < 0·0001). Similarly, we found higher pooled prevalence estimates for studies from Turkey (76·3%, 95% CI 60·9-87·0; I^2^= 91·1%, 95% CI 84·2-95·0, p < 0·0001). We also grouped studies by geography and economic development. Group 1: USA, Chile, Australia, Canada, Ireland, Japan, Spain; group 2: South Africa, China, Egypt, Iran, Turkey, Saudi Arabia; and group 3: Bangladesh, Thailand, and India. Prevalence was 36·1% (95% CI 27·8-45·4) for group 1 (n = 18), 62·7% (95% CI 52·2-72·2) for group 2 (n = 18) and 71·4% (95% CI 57·9-82·0) for group 3 (n = 12) *(Additional Figure 2).* Variation attributable to heterogeneity was still high in the three subgroups (I^2^> 90%).

Given the broad age range in included studies, we combined studies with only neonates^48, 60, 62-65^ and observed a prevalence estimate of 85·6% (95% CI 78·5-90·6) with moderate variation attributable to heterogeneity (I^2^= 54·3%, 95% CI 0·0-81·7, p value = 0·05). In all other studies (n = 42) that included children of broad age ranges, estimated prevalence was lower at 49·7% (95% CI 42·9-56·5; I^2^= 94·7%, 95% CI 93·6-95·6, p value < 0·0001) (*Additional Table 13, Additional Figure 3*).

### Post-hoc investigation to determine sources of heterogeneity

To investigate the substantial heterogeneity observed in prevalence estimates, we incorporated study-specific characteristics (year of publication, total study sample size, quality score, study design, country group and clinical setting) as covariates in a random effects meta-regression model. We identified clinical setting and country groups as significant predictors *(Figure 3).* We found that the model fitted with all available covariates can explain 37·52% of I^2^ with F = 5·1119, p = 0·0005 *(Additional Table 14).* We also conducted univariate meta-regressions for each of the six predictors *(Additional Figure 4).*

**Figure 3.**
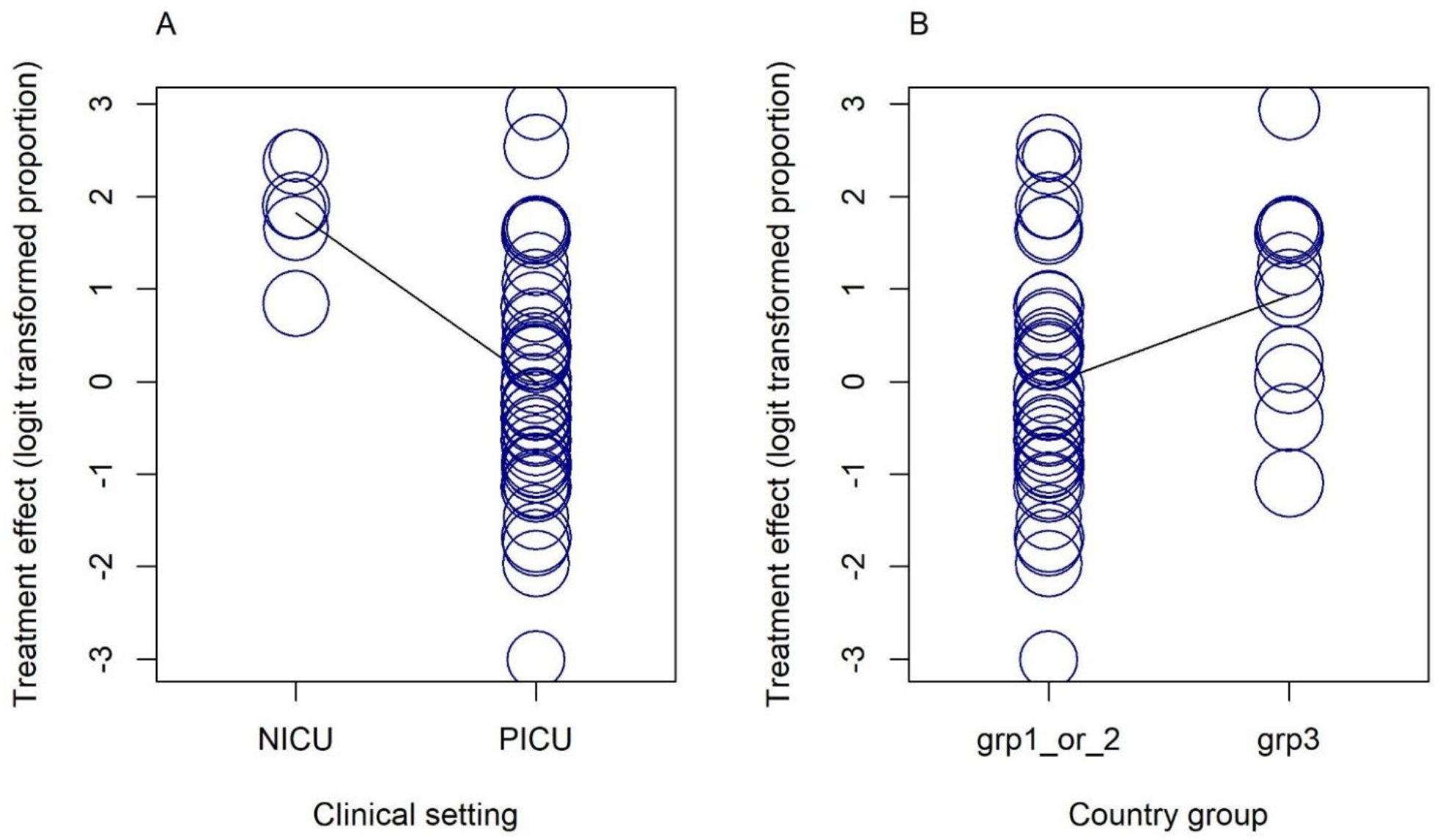
Bubble plots of univariate meta-regressions. Each study is represented by a circle. Predictor variables; A clinical setting and B country groups are shown on the x-axis and the effect measure logit transformed proportion shown on the vertical (y-axis). NICU = Neonatal Intensive Care Unit; PICU = Pediatric Intensive Care Unit; grp = country group; country group 1 = USA, Chile, Australia, Canada, Ireland, Japan, Spain; country group 2 = South Africa, China, Egypt, Iran, Turkey, Saudi Arabia; and country group 3 = Bangladesh, Thailand, and India

### Prevalence of vitamin D deficiency in critically ill children with sepsis and in those with respiratory tract infections

A total of 788 (median 42, range 9 -160) patients had a diagnosis of sepsis, of which 499 (63·3%) were vitamin D deficient. Nine of the sixteen studies including septic patients were cohort (56·3%) and seven (43·8%) case-control *(Additional Table 15).* Most studies originated from India (n = 6) Turkey (n = 3) or Ireland (n = 2) and 15 were published between 2014 and 2017. Thirteen studies took place in a PICU and the remaining^60, 63, 65^ in NICUs. We found that all studies were of medium to high quality (median NOS score 6·5, range 5 – 8). Pooled prevalence of vitamin D deficiency was 63·8% (95% CI 49·9-75·7) *(Figure 4)*. Variation attributable to heterogeneity was high (I^2^= 90·5%, 95% CI 86·2-93·5%, p < 0·0001). Funnel plot was symmetric suggesting no publication bias (p = 0·828, Egger’s test) *(Additional Figure 5).*

**Figure 4.**
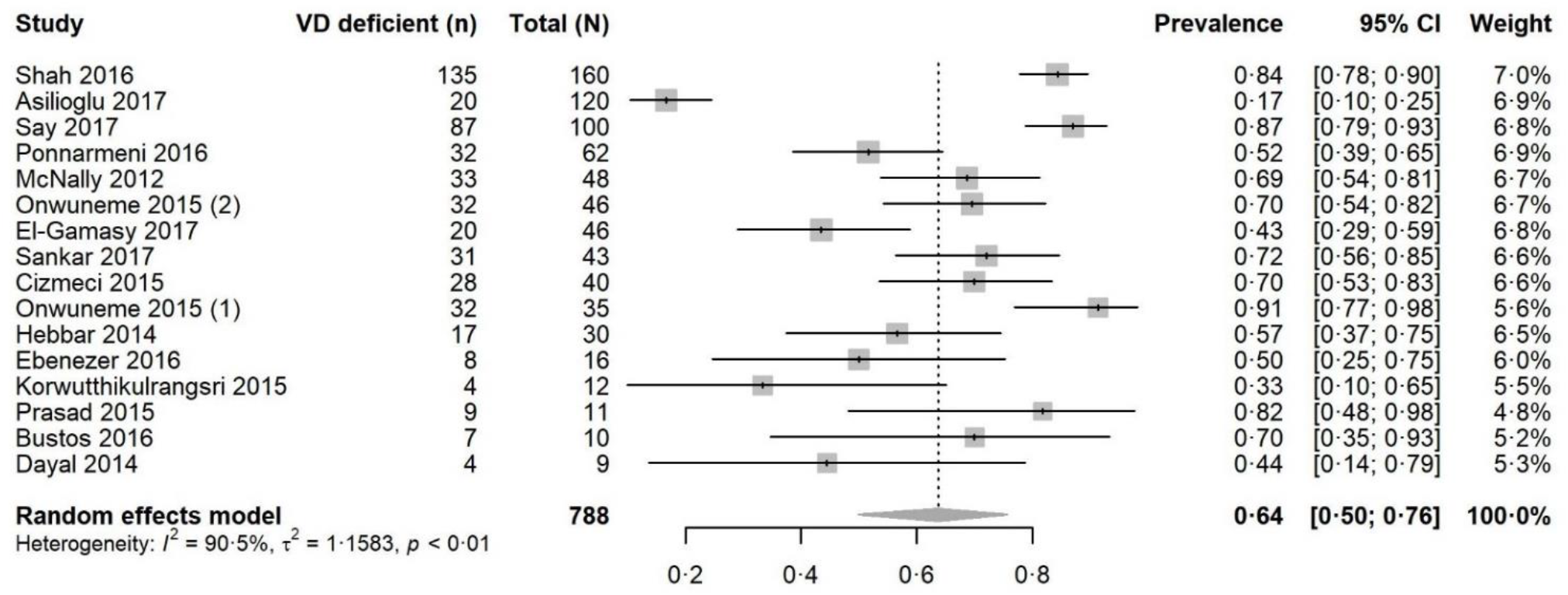
Pooled prevalence estimate for vitamin D deficiency in critically ill children with sepsis. Forest plot shows result from the random effects model. The diamond represents the overall pooled proportion of vitamin D deficiency from the meta-analysis of the 16 studies. Each square shows the prevalence estimate of each study and the horizontal line across each square represents the 95% confidence interval (CI) of the prevalence estimate.

We also analysed studies of patients admitted for respiratory tract infections (n = 24) such as acute lower respiratory tract infection (ALRTI), pneumonia and bronchiolitis. Of these 1,683 total individuals (median 49), 778 (46·2%) were vitamin D deficient. These studies were of high to medium quality (median NOS score 6·5, range 6 - 8). Most originated from India (n = 6) and Spain (n = 4). We found a prevalence estimate of 49·9% (95% CI 37·6-62·2; I^2^= 93·9%, 95% CI 92·1-95·3, p < 0·0001), with no evidence of publication bias (p = 0·217, Egger’s test) (*Table 1)*. Two of these studies^50, 76^ also investigated sepsis.

### Sensitivity analysis for prevalence in children with sepsis

Exclusion of the studies^58, 60, 65, 77^ utilising thresholds other than < 50 nmol/L for deficiency yielded a similar estimate of prevalence at 61·4% (95% CI 43·5-76·6; I^2^= 91·2% 86·5-94·2, p < 0·0001) *(Additional Table 16).*

We examined pooled prevalence estimates according to sample size (< 40 versus ≥ 40). Studies with a small sample size (n = 7; 123 total individuals) showed a prevalence estimate of 63·2% (95% CI 45·0-78·2) with moderate variation attributable to heterogeneity (I^2^= 66·2%, 95% CI 24·5-84·9, p = 0·0068). For the remaining nine studies (sample sizes ≥ 40, 665 total individuals) the estimate was 63·9% (95% CI 44·9 - 79·4) with high variation attributable to heterogeneity (I^2^= 94·3%, 95% CI 91·2-96·3, p < 0·0001).

There was no material change in prevalence estimates when analysed according to study design. The nine cohort studies (463 total individuals) gave an estimate of 62·6% (95% CI 40·7-80·4) with high variation attributable to heterogeneity (I^2^= 92·8%, 95% CI 88·6-95·5, p < 0·0001). Case-control studies (n = 7; 325 total individuals) showed a prevalence of 65·2% (95% CI 47·3-79·7; I^2^= 87·0%, 95% CI 75·5-93·1, p < 0·0001) *(Additional Table 16, Additional Figure 6).*

Studies from India (n = 6) gave a prevalence estimate of 66·4% (95% CI 48·3-80·7; I^2^= 83·6%, 95% CI 65·7-92·2, < 0·0001). The three studies from Turkey assessing septic patients gave a pooled estimate of 59·2% (95% CI 13·6-93·1; I^2^= 97·8%, 95% CI 95·8-98·8, p < 0·0001) *(Additional Table 16).*

The prevalence estimate in the three studies^60, 63, 65^ including neonates with sepsis was 76·9% (95% CI 61·9-87·3, I^2^= 74·7%, 95% CI 15·9-92·4, p-value 0·019). The thirteen studies with children of different ages, excluding neonates, gave a pooled estimate of 60·1% (95% CI 43·7-74·5; I^2^= 90·8%, 95% CI 86·1-93·9, p value < 0·0001) *(Additional Table 16).*

### Mortality in critically ill children

We identified 18 studies^8, 47, 50-53, 55, 58, 59, 65, 68, 69, 71, 76-80^ assessing vitamin D status and mortality. These studies included a total of 2,463 individuals, from which 220 deaths (17·2%) were observed in 1,278 (51·9%) individuals with vitamin D deficiency and 99 deaths (8·4%) were observed in 1,185 individuals without deficiency (48·1%).

All 18 studies took place in a PICU apart from one^65^ in a NICU. Sixteen of these studies (89%) were published between 2014 and 2017. Fourteen were cohort (77·8%) and four case-controls (22·2%). Almost half (n = 7) of the studies originated from India. Quality scores ranged from 5 to 8 with a median of 6.

Using a random effects model, we found that vitamin D deficiency in critically ill children significantly increased the risk of death (OR 1·81, 95% CI 1·24-2·64, p-value = 0·002) with low, non-significant heterogeneity (I^2^= 25·7%, 95% CI 0·0-58·0, p = 0·153) (*Table 1, Figure 5*). We did not identify evidence of publication bias (p = 0·084, Egger’s test) (*Additional Figure 7*).

**Figure 5.**
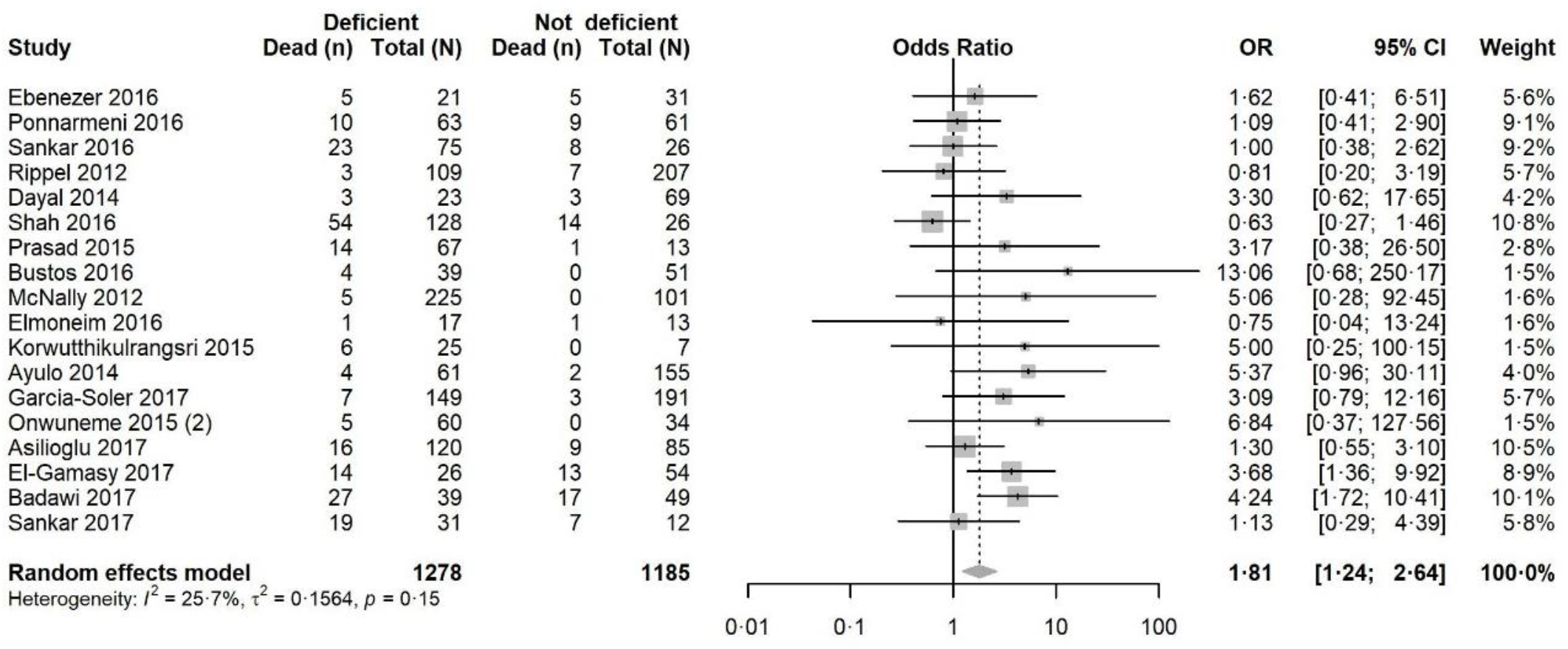
Pooled odds ratio (OR) of risk of mortality in vitamin D deficient versus vitamin D non-deficient critically ill children. Forest plot shows result from the random effects model. Diamond represents the overall OR (with corresponding 95% Confidence Interval). Each square shows the odds ratio of each study and the horizontal line across each square represents the 95% confidence interval (CI) of the estimate.

### Sensitivity analysis for mortality in critically ill children

We obtained similar results through the fixed effects model (OR 1·72, 95% CI 1·27-2·33, p = 0·0005) (*Table 1, Additional Figure 8*). When excluding studies with thresholds other than < 50 nmol/L indicating deficiency, we found the association between vitamin D deficiency and increased risk of mortality still significant but lower, both with the random (OR 1·59, 95% CI 1·05-2·41, p = 0·028; I^2^= 24·3%, 95% CI 0·0-59·9, p = 0·191) and fixed effect models (OR 1·52, 95% CI 1·08-2·13, p = 0·016) with no indication of publication bias (p = 0·12, Egger’s test) *(Additional Table 17).*

A significant association was also observed in analysis of the 14 cohort studies, both with the random (OR 1·80, 95% CI 1·15-2·81, p = 0·01) and fixed effects model (OR 1·65, 95% CI 1·17-2·34, p-value = 0·004) with low variation attributable to heterogeneity (I^2^= 31·3%, 95% CI 0·0-63·7). Pooling the four case-control studies together we obtained a significant positive association with the fixed (OR 1·97, 95% CI 1·02-3·82, p = 0·044) effects model but non-significant with the random effects model (OR 1·97, 95% CI 0·88-4·42, p = 0·098). The association was positive but not-significant when pooling the seven studies from India with the random effects model (OR 1·08, 95% CI 0·70-1·69, p-value = 0·710; I^2^= 0·0% 0·0-62·4, p = 0·589) and similar with fixed effects (OR 1·08, 95% CI 0·70-1·69, p = 0·710) *(Additional Table 16)*.

### Mortality in patients with sepsis and respiratory tract infections

We were unable to identify a sufficient number of studies assessing vitamin D and mortality for meta-analysis in individuals with sepsis. Three studies^8, 58, 60^ measured vitamin D levels in paediatric patients with sepsis. One study^8^ assessed mortality and did not find a significant association in children from 1 to 12 years with sepsis (n=124). None of the studies with children admitted for respiratory tract infections looked at the association of vitamin D deficient versus vitamin D not deficient children with mortality.

## Discussion

Vitamin D deficiency is highly prevalent worldwide, even in countries with abundant sunshine. Studies demonstrated high prevalence of vitamin D deficiency in otherwise healthy children from high-income countries (9 to 24%) but also from middle and low-income countries in Indian subcontinent (36 to 90%).^8^

We identified 48 studies representing a total of 5,520 children treated in ICU or emergency units for acute conditions who had blood vitamin D levels measured close to or upon admission. Our analysis shows that prevalence of vitamin D deficiency is high (range 5%^66^ to 95%^54^) across ICU and emergency units in the paediatric population, particularly in individuals with sepsis. Importantly, our analysis showed a significantly increased risk of mortality in critically ill children with vitamin D deficiency. We carried out several analyses for sensitivity including fixed effects models, by study design, country group, age and sample size and found consistent results. A recently published meta-analysis^81^ also investigated prevalence of vitamin D deficiency in critically ill children and its association with risk of mortality and showed similar results to ours.

Sub-group analyses in patients with sepsis or respiratory tract infections demonstrated a high prevalence of vitamin D deficiency, consistent with the increased risk of bacterial or nosocomial infection in vitamin D deficient individuals identified elsewhere.^81^

Although sepsis is a leading cause of paediatric mortality and morbidity worldwide,^82^ we found few studies assessing the relationship between vitamin D status and mortality in this population. We were unable to identify sufficient studies including patients with sepsis to perform a meta-analysis of vitamin D status and mortality. Sepsis remains an area of unmet need with high social and financial costs.^24^ Diagnostic criteria,^83^ a lack of adequate biomarkers^84^ and targeted treatment remain important challenges in research on sepsis. We did not find studies that assessed the risk of mortality in relation to vitamin D deficiency in children admitted for respiratory tract infections either.

Strengths of our review include the large number of studies and large total sample size, allowing a high-powered investigation to identify meaningful associations. For our systematic review and meta-analysis, we followed pre-specified eligibility criteria and used the PRISMA^27^ and MOOSE guidelines^28^ for reporting. We carried out sensitivity analyses with few material differences in results. However, we note that the relationship between vitamin D deficiency and mortality was sensitive to study design and studies from India, probably due to the smaller number of individuals in those analyses. Only the prevalence analysis with neonates indicated lower variation attributable to heterogeneity (I^2^= 54·3%) along with a higher prevalence estimate (86%) compared to other analyses. As expected, heterogeneity across studies is high overall, particularly for prevalence estimates. We utilised meta-regression to investigate this substantial heterogeneity around prevalence estimates. From the six variables in our multi-variable model, only clinical setting and country groups were found to be significant predictors of pooled prevalence estimates of vitamin D deficiency and the full model could explain 37·52% of I^2^. Studies in NICU yielded higher prevalence estimates compared to studies in PICU. Studies from group 3 countries were also associated with higher prevalence estimates compared to studies from countries of group 1 and 2. Other variables, mainly individual patient characteristics such as age and ethnicity, were not directly available to us and may account for significant heterogeneity. Future research should also investigate biological heterogeneity in order to strengthen the evidence and produce generalisable results.

Our systematic review did not identify longitudinal studies with multiple time-point, pre-disease or pre-admission vitamin D measurements. The majority of studies were single centre with heterogeneous patient groups and relatively small sample sizes. Few studies accounted for important confounders that influence vitamin D levels such as age, gender, BMI, season of measurements, vitamin D supplementation and comorbidities. The relationship observed between vitamin D deficiency and mortality could be due to reverse causation and future studies will need to control for these covariates and other confounders.

Although included studies were generally of good quality, sample sizes varied considerably and were typically small. Over half of studies included less than 100 cases and only 10 studies (19·6%) had a total sample size of more than 200 individuals. In addition, studies used a variety of definitions and age ranges to designate individuals as children. Our analysis only included mortality as a clinical outcome. A further general limitation is the difference in thresholds for vitamin D deficiency, particularly in the levels which are considered normal for infants and young children. Our assessment used the currently recommended threshold for deficiency (≤ 50 nmol/L)^16^ and used a conservative estimate for studies which used different criteria.

Vitamin D remains an attractive biomarker and potential therapeutic agent in acute and critical care patients. Carefully designed and adequately powered studies are needed to determine the importance and therapeutic value of vitamin D in the general and septic paediatric critical care population.

## Availability of data and materials

Data and computational code used for processing and analysis are available at https://github.com/margarc/VitaminD_children

## Author contributions

AJBT conceived the study. AJBT and IT designed the study. MC collected data and performed the analysis with input from MAC, IT, ABJT and EE. MC and AJBT wrote the manuscript with contributions from all authors.

## Declaration of interests

The authors declare no conflicts of interest.

## Acknowledgements

AJBT was supported by the Medical Research Council (UK MED-BIO Programme Fellowship, MR/L01632X/1).

## Ethics committee approval

Not applicable.

## Supplementary Appendix

### Additional Tables

Additional Table 1A Search terms used in OVID

Additional Table 1B Search terms used in PubMed

Additional Table 2A Newcastle Ottawa study quality scoring system (cohort studies)

Additional Table 2B Newcastle Ottawa study quality scoring system (case-control studies)

Additional Table 2C Newcastle Ottawa study quality scoring system (cross sectional studies)

Additional Table 3A Circulating 25(OH) D threshold levels used in the selected studies

Additional Table 3B Circulating 25(OH) D threshold levels used in the selected studies for prevalence in sepsis

Additional Table 4 Studies with thresholds other than <50 nmol/L

Additional Table 5 Excluded studies

Additional Table 6 Characteristics of the 48 included studies

Additional Table 7 Objectives and outcomes of included studies

Additional Table 8 Assay used in each study to measure Vitamin D levels

Additional Table 9 Funding and ethical approval of included studies

Additional Table 10 Age groups of children in each study

Additional Table 11 Prevalence of vitamin D deficiency in each study of critically ill children (sorted from highest to lowest)

Additional Table 12 Characteristics of studies used in the meta-analysis for prevalence

Additional Table 13 Sensitivity analyses for prevalence of vitamin D deficiency in all critically ill children

Additional Table 14 Multivariate meta-regression model for prevalence

Additional Table 15 Characteristics of studies included in the meta-analysis for prevalence in individuals with sepsis

Additional Table 16 Sensitivity analyses for prevalence of vitamin D deficiency in critically ill children with sepsis

Additional Table 17 Sensitivity analyses for mortality

### Additional Figures

Additional Figure 1 Funnel plot of studies of prevalence of vitamin D deficiency in critically ill children

Additional Figure 2 Pooled prevalence estimate for vitamin D deficiency in critically ill children (subgroup analysis by country group)

Additional Figure 3 Pooled prevalence estimate for vitamin D deficiency in critically ill children (subgroup analysis of neonates versus all other age groups)

Additional Figure 4 Bubble plots of univariate meta-regressions.

Additional Figure 5 Funnel plot for prevalence of vitamin D deficiency in critically ill children with sepsis

Additional Figure 6 Pooled prevalence estimate for vitamin D deficiency in critically ill children with sepsis (subgroup analysis by study design)

Additional Figure 7 Funnel plot of risk of mortality in vitamin D deficient versus vitamin D non-deficient critically ill children

Additional Figure 8 Pooled odds ratio and 95% CI for risk of mortality in vitamin D deficient versus vitamin D non-deficient critically ill children (fixed effects model)

## References

1. Shuler FD, Wingate MK, Moore GH, Giangarra C. Sports health benefits of vitamin d. Sports health. 2012;4(6):496–501.

2. Verceles AC, Weiler B, Koldobskiy D, Goldberg AP, Netzer G, Sorkin JD. Association Between Vitamin D Status and Weaning From Prolonged Mechanical Ventilation in Survivors of Critical Illness. Respiratory care. 2015;60(7):1033–9.

3. Braegger C, Campoy C, Colomb V, Decsi T, Domellof M, Fewtrell M, et al Vitamin D in the healthy European paediatric population. Journal of pediatric gastroenterology and nutrition. 2013;56(6):692–701.

4. Holick MF, Binkley NC, Bischoff-Ferrari HA, Gordon CM, Hanley DA, Heaney RP, et al Evaluation, treatment, and prevention of vitamin D deficiency: an Endocrine Society clinical practice guideline. The Journal of clinical endocrinology and metabolism. 2011;96(7):1911–30.

5. Holick MF. Vitamin D deficiency. The New England journal of medicine. 2007;357(3):266–81.

6. Aranow C. Vitamin D and the immune system. Journal of investigative medicine: the official publication of the American Federation for Clinical Research. 2011;59(6):881–6.

7. Zhang YP, Wan YD, Sun TW, Kan QC, Wang LX. Association between vitamin D deficiency and mortality in critically ill adult patients: a meta-analysis of cohort studies. Critical care (London, England). 2014;18(6):684.

8. Ponnarmeni S, Kumar Angurana S, Singhi S, Bansal A, Dayal D, Kaur R, et al Vitamin D deficiency in critically ill children with sepsis. Paediatrics and international child health. 2016;36(1):15–21.

9. Kulie T, Groff A, Redmer J, Hounshell J, Schrager S. Vitamin D: an evidence-based review. Journal of the American Board of Family Medicine: JABFM. 2009;22(6):698–706.

10. Hewison M. Antibacterial effects of vitamin D. Nature reviews Endocrinology. 2011;7(6):337–45.

11. Kempker JA, West KG, Kempker RR, Siwamogsatham O, Alvarez JA, Tangpricha V, et al Vitamin D status and the risk for hospital-acquired infections in critically ill adults: a prospective cohort study. PLoS one. 2015;10(4):e0122136.

12. Martineau AR, Jolliffe DA, Hooper RL, Greenberg L, Aloia JF, Bergman P, et al Vitamin D supplementation to prevent acute respiratory tract infections: systematic review and meta-analysis of individual participant data. BMJ (Clinical research ed). 2017;356:i6583.

13. Absoud M, Cummins C, Lim MJ, Wassmer E, Shaw N. Prevalence and predictors of vitamin D insufficiency in children: a Great Britain population based study. PLoS one. 2011;6(7):e22179.

14. Greer FR. Defining vitamin D deficiency in children: beyond 25-OH vitamin D serum concentrations. Pediatrics. 2009;124(5):1471–3.

15. Abou-Zahr R, Kandil SB. A pediatric critical care perspective on vitamin D. Pediatric research. 2015;77(1-2):164–7.

16. Ross AC, Manson JE, Abrams SA, Aloia JF, Brannon PM, Clinton SK, et al The 2011 report on dietary reference intakes for calcium and vitamin D from the Institute of Medicine: what clinicians need to know. The Journal of clinical endocrinology and metabolism. 2011;96(1):53–8.

17. Ala-Kokko TI, Mutt SJ, Nisula S, Koskenkari J, Liisanantti J, Ohtonen P, et al Vitamin D deficiency at admission is not associated with 90-day mortality in patients with severe sepsis or septic shock: Observational FINNAKI cohort study. Annals of medicine. 2016;48(1-2):67–75.

18. Putzu A, Belletti A, Cassina T, Clivio S, Monti G, Zangrillo A, et al Vitamin D and outcomes in adult critically ill patients. A systematic review and meta-analysis of randomized trials. Journal of critical care. 2017;38:109–14.

19. Han JE, Jones JL, Tangpricha V, Brown MA, Brown LAS, Hao L, et al High Dose Vitamin D Administration in Ventilated Intensive Care Unit Patients: A Pilot Double Blind Randomized Controlled Trial. Journal of clinical & translational endocrinology. 2016;4:59–65.

20. Quraishi SA, De Pascale G, Needleman JS, Nakazawa H, Kaneki M, Bajwa EK, et al Effect of Cholecalciferol Supplementation on Vitamin D Status and Cathelicidin Levels in Sepsis: A Randomized, Placebo-Controlled Trial. Critical care medicine. 2015;43(9):1928–37.

21. Singer M, Deutschman CS, Seymour CW, Shankar-Hari M, Annane D, Bauer M, et al The Third International Consensus Definitions for Sepsis and Septic Shock (Sepsis-3). Jama. 2016;315(8):801–10.

22. NHS-England. Improving outcomes for patients with sepsis: a cross-system action plan 2015 [Available from: https://www.england.nhs.uk/ourwork/part-rel/sepsis/.

23. Plunkett A, Tong J. Sepsis in children. BMJ (Clinical research ed). 2015;350:h3017.

24. Weiss SL, Fitzgerald JC, Pappachan J, Wheeler D, Jaramillo-Bustamante JC, Salloo A, et al Global epidemiology of pediatric severe sepsis: the sepsis prevalence, outcomes, and therapies study. American journal of respiratory and critical care medicine. 2015;191(10):1147–57.

25. Mayr FB, Yende S, Angus DC. Epidemiology of severe sepsis. Virulence. 2014;5(1):4–11.

26. De Pascale G, Ranzani OT, Nseir S, Chastre J, Welte T, Antonelli M, et al Intensive care unit patients with lower respiratory tract nosocomial infections: the ENIRRIs project. ERJ open research. 2017;3(4).

27. Moher D, Liberati A, Tetzlaff J, Altman DG. Preferred reporting items for systematic reviews and meta-analyses: the PRISMA statement. Journal of clinical epidemiology. 2009;62(10):1006–12.

28. Stroup DF, Berlin JA, Morton SC, Olkin I, Williamson GD, Rennie D, et al Meta-analysis of observational studies in epidemiology: a proposal for reporting. Meta-analysis Of Observational Studies in Epidemiology (MOOSE) group. Jama. 2000;283(15):2008–12.

29. Wells GA SB OCD, Peterson J, Welch V, Losos M,Tugwell P. The Newcastle-Ottawa Scale (NOS) for assessing the quality of nonrandomised studies in meta-analyses [Available from: http://www.ohri.ca/programs/clinical_epidemiology/oxford.asp.

30. Newcombe RG. Two-sided confidence intervals for the single proportion: comparison of seven methods. Statistics in medicine. 1998;17(8):857–72.

31. DerSimonian R, Laird N. Meta-analysis in clinical trials revisited. Contemporary clinical trials. 2015;45(Pt A):139–45.

32. Borenstein M, Hedges LV, Higgins JP, Rothstein HR. A basic introduction to fixed-effect and random-effects models for meta-analysis. Research synthesis methods. 2010;1(2):97–111.

33. Higgins JP, Thompson SG, Deeks JJ, Altman DG. Measuring inconsistency in meta-analyses. BMJ (Clinical research ed). 2003;327(7414):557–60.

34. Higgins JP, Thompson SG. Quantifying heterogeneity in a meta-analysis. Statistics in medicine. 2002;21(11):1539–58.

35. Egger M, Davey Smith G, Schneider M, Minder C. Bias in meta-analysis detected by a simple, graphical test. BMJ (Clinical research ed). 1997;315(7109):629–34.

36. Panityakul T, Bumrungsup C, Knapp G. On estimating residual heterogeneity in random-effects meta-regression: a comparative study. J Stat Theory Appl. 2013;12(3):253.

37. Knapp G, Hartung J. Improved tests for a random effects meta-regression with a single covariate. Statistics in medicine. 2003;22(17):2693–710.

38. Schwarzer G. meta: An R package for meta-analysis. R news. 2007;7(3):40–5.

39. Viechtbauer W. Conducting meta-analyses in R with the metafor package. Journal of statistical software. 2010;36(3).

40. Garg D, Sharma VK, Karnawat B. Association of serum vitamin D with acute lower respiratory infection in Indian children under 5 years: a case control study. International Journal of Contemporary Pediatrics. 2016;3(4):1164–9.

41. Aydemir G, Cekmez F, Kalkan G, Fidanci MK, Kaya G, Karaoglu A, et al High serum 25-hydroxyvitamin D levels are associated with pediatric sepsis. The Tohoku journal of experimental medicine. 2014;234(4):295–8.

42. Gamal TS, Madiha AS, Hanan MK, Abdel-Azeem ME, Marian GS. Neonatal and Maternal 25-OH Vitamin D Serum Levels in Neonates with Early-Onset Sepsis. Children (Basel, Switzerland). 2017;4(5).

43. Seliem MS, Abdel Haie OM, Mansour AI, Salama SSME. The relation between vitamin D level and increased risk for early-onset neonatal sepsis in full-term infants. Medical Research Journal. 2016;15(1):16–21.

44. McNally JD, Leis K, Matheson LA, Karuananyake C, Sankaran K, Rosenberg AM. Vitamin D deficiency in young children with severe acute lower respiratory infection. Pediatric pulmonology. 2009;44(10):981–8.

45. McNally JD, Doherty DR, Lawson ML, Al-Dirbashi OY, Chakraborty P, Ramsay T, et al The relationship between vitamin D status and adrenal insufficiency in critically ill children. The Journal of clinical endocrinology and metabolism. 2013;98(5):E877–81.

46. McNally JD, Iliriani K, Pojsupap S, Sampson M, O’Hearn K, McIntyre L, et al Rapid normalization of vitamin D levels: a meta-analysis. Pediatrics. 2015;135(1):e152–66.

47. McNally JD, Menon K, Chakraborty P, Fisher L, Williams KA, Al-Dirbashi OY, et al The association of vitamin D status with pediatric critical illness.Pediatrics. 2012;130(3):429–36.

48. Karatekin G, Kaya A, Salihoglu O, Balci H, Nuhoglu A. Association of subclinical vitamin D deficiency in newborns with acute lower respiratory infection and their mothers. European journal of clinical nutrition. 2009;63(4):473–7.

49. Madden K, Feldman HA, Smith EM, Gordon CM, Keisling SM, Sullivan RM, et al Vitamin D deficiency in critically ill children. Pediatrics. 2012;130(3):421–8.

50. Shah SK, Kabra SK, Gupta N, Pai G, Lodha R. Vitamin D Deficiency and Parathyroid Response in Critically-ill Children: Association with Illness Severity and Clinical Outcomes. Indian pediatrics. 2016;53(6):479–84.

51. Sankar J, Lotha W, Ismail J, Anubhuti C, Meena RS, Sankar MJ. Vitamin D deficiency and length of pediatric intensive care unit stay: a prospective observational study. Annals of intensive care. 2016;6(1):3.

52. Dayal D, Kumar S, Sachdeva N, Kumar R, Singh M, Singhi S. Fall in Vitamin D Levels during Hospitalization in Children. International journal of pediatrics. 2014;2014:291856.

53. Prasad S, Raj D, Warsi S, Chowdhary S. Vitamin D Deficiency and Critical Illness. Indian journal of pediatrics. 2015;82(11):991–5.

54. Wayse V, Yousafzai A, Mogale K, Filteau S. Association of subclinical vitamin D deficiency with severe acute lower respiratory infection in Indian children under 5 y. European journal of clinical nutrition. 2004;58(4):563–7.

55. Ebenezer K, Job V, Antonisamy B, Dawodu A, Manivachagan MN, Steinhoff M. Serum Vitamin D Status and Outcome among Critically Ill Children Admitted to the Pediatric Intensive Care Unit in South India. Indian journal of pediatrics. 2016;83(2):120–5.

56. Narang GS, Arora S, Kukreja S. Association of Vitamin D Deficiency with Acute Lower Respiratory Infection in Toddlers. Journal of Nepal Paediatric Society. 2016;36(1):14–6.

57. Jat KR, Kaur J, Guglani V. Vitamin D and Pneumonia in Children: A Case Control Study. J Pulm Med Respir Res. 2016;2(004).

58. Sankar J, Ismail J, Das R, Dev N, Chitkara A, Sankar MJ. Effect of Severe Vitamin D Deficiency at Admission on Shock Reversal in Children With Septic Shock. Journal of intensive care medicine. 2017:885066617699802.

59. Asilioglu N, Cigdem H, Paksu MS. Serum Vitamin D Status and Outcome in Critically Ill Children. Indian journal of critical care medicine: peer-reviewed, official publication of Indian Society of Critical Care Medicine. 2017;21(10):660–4.

60. Say B, Uras N, Sahin S, Degirmencioglu H, Oguz SS, Canpolat FE. Effects of cord blood vitamin D levels on the risk of neonatal sepsis in premature infants. Korean journal of pediatrics. 2017;60(8):248– 53.

61. Cayir A, Turan MI, Ozkan O, Cayir Y. Vitamin D levels in children diagnosed with acute otitis media. JPMA The Journal of the Pakistan Medical Association. 2014;64(11):1274–7.

62. Cetinkaya M, Cekmez F, Buyukkale G, Erener-Ercan T, Demir F, Tunc T, et al Lower vitamin D levels are associated with increased risk of early-onset neonatal sepsis in term infants. Journal of Perinatology. 2015;35(1):39.

63. Cizmeci MN, Kanburoglu MK, Akelma AZ, Ayyildiz A, Kutukoglu I, Malli DD, et al Cord-blood 25-hydroxyvitamin D levels and risk of early-onset neonatal sepsis: a case-control study from a tertiary care center in Turkey. European journal of pediatrics. 2015;174(6):809–15.

64. Dinlen N, Zenciroglu A, Beken S, Dursun A, Dilli D, Okumus N. Association of vitamin D deficiency with acute lower respiratory tract infections in newborns. The Journal of maternal-fetal & neonatal medicine: the official journal of the European Association of Perinatal Medicine, the Federation of Asia and Oceania Perinatal Societies, the International Society of Perinatal Obstet. 2016;29(6):928–32.

65. Onwuneme C, Martin F, McCarthy R, Carroll A, Segurado R, Murphy J, et al The Association of Vitamin D Status with Acute Respiratory Morbidity in Preterm Infants. The Journal of pediatrics. 2015;166(5):1175-80.e1.

66. Roth DE, Jones AB, Prosser C, Robinson JL, Vohra S. Vitamin D status is not associated with the risk of hospitalization for acute bronchiolitis in early childhood. European journal of clinical nutrition. 2009;63(2):297–9.

67. Cebey-Lopez M, Pardo-Seco J, Gomez-Carballa A, Martinon-Torres N, Rivero-Calle I, Justicia A, et al Role of Vitamin D in Hospitalized Children With Lower Tract Acute Respiratory Infections. Journal of pediatric gastroenterology and nutrition. 2016;62(3):479–85.

68. García-Soler P, Morales-Martínez A, Rosa-Camacho V, Lillo-Muñoz JA, Milano-Manso G. Vitamin D deficiency and morbimortality in critically ill paediatric patients. Anales de Pediatría (English Edition). 2017;87(2):95–103.

69. Rippel C, South M, Butt WW, Shekerdemian LS. Vitamin D status in critically ill children. Intensive care medicine. 2012;38(12):2055–62.

70. Alonso MA, Pallavicini ZF, Rodriguez J, Avello N, Martinez-Camblor P, Santos F. Can vitamin D status be assessed by serum 25OHD in children? Pediatric nephrology (Berlin, Germany). 2015;30(2):327–32.

71. Ayulo M, Jr., Katyal C, Agarwal C, Sweberg T, Rastogi D, Markowitz M, et al The prevalence of vitamin D deficiency and its relationship with disease severity in an urban pediatric critical care unit. Endocrine regulations. 2014;48(2):69–76.

72. Rey C, Sanchez-Arango D, Lopez-Herce J, Martinez-Camblor P, Garcia-Hernandez I, Prieto B, et al Vitamin D deficiency at pediatric intensive care admission. Jornal de pediatria. 2014;90(2):135–42.

73. Onwuneme C, Carroll A, Doherty D, Bruell H, Segurado R, Kilbane M, et al Inadequate vitamin D levels are associated with culture positive sepsis and poor outcomes in paediatric intensive care. Acta paediatrica (Oslo, Norway: 1992). 2015;104(10):e433–8.

74. Jia K-P, Zhao L-F, Feng N, Ma K, Li Y-X. Lower level of vitamin D3 is associated with susceptibility to acute lower respiratory tract infection (ALRTI) and severity: a hospital based study in Chinese infants. INTERNATIONAL JOURNAL OF CLINICAL AND EXPERIMENTAL MEDICINE. 2017;10(5):7997–8003.

75. Halwany AS, Deghady AAE, Moustafa AAA, Diab AEM. Study of Serum Vitamin D Level among Critically Ill Patients Admitted to Alexandria University Pediatric Intensive Care Unit. 2017.

76. Bustos R, Rodriguez-Nunez I, Peña RZ, Soto GG. Vitamin D deficiency in children admitted to the paediatric intensive care unit. Revista chilena de pediatria. 2016;87(6):480–6.

77. El-Gamasy M, Eldeeb M, Abdelmageed M. Prognostic Value of Vitamin D Status in Pediatric Patients with Acute Kidney Injury in Tanta University Emergency Hospital, Egypt. J Emerg Intern Med. 2017;1(1):5.

78. Elmoneim A, Rhill R, Rahalli M, Al-Rahalli M, Alrehally A, Alrohily S. Vitamin D level in pediatric intensive care unit (PICU) patients: its relation to severity of illness. Pediat Therapeut. 2016;6(293):2161-0665.1000293.

79. Korwutthikulrangsri M, Mahachoklertwattana P, Lertbunrian R, Chailurkit LO, Poomthavorn P. Vitamin D deficiency and adrenal function in critically ill children. Journal of the Medical Association of Thailand = Chotmaihet thangphaet. 2015;98(4):365–72.

80. Badawi NES, Algebaly HAF, El Sayed R, Zeid ESA. Vitamin D deficiency in critically ill children. Kasr Al Ainy Medical Journal. 2017;23(1):6.

81. McNally JD, Nama N, O’Hearn K, Sampson M, Amrein K, Iliriani K, et al Vitamin D deficiency in critically ill children: a systematic review and meta-analysis. Critical care (London, England). 2017;21(1):287.

82. Randolph AG, McCulloh RJ. Pediatric sepsis: important considerations for diagnosing and managing severe infections in infants, children, and adolescents. Virulence. 2014;5(1):179–89.

83. Vincent JL. The Clinical Challenge of Sepsis Identification and Monitoring. PLoS medicine. 2016;13(5):e1002022.

84. Mathias B, Mira JC, Larson SD. Pediatric sepsis. Current opinion in pediatrics. 2016;28(3):380–7.

